# Trans-generational effect of protein restricted diet on adult body and wing size of *Drosophila melanogaster*

**DOI:** 10.1101/2020.02.17.952101

**Authors:** Sudhakar Krittika, Pankaj Yadav

## Abstract

Dietary Restriction (DR) via protein restriction (PR) has become an inquisitive field and has established feasible trade-offs between various fitness and behavioral traits in *Drosophila melanogaster* to understand lifespan or aging in a nutritionally challenged environment. However, the phenotypes of body size, weight and wing length respond according to factors such as flies’ genotype, environmental exposure, and parental diet. Hence, understanding the long-term effect of PR on these phenotypes is essential. Here, we demonstrate the effect of PR diet on body size, weight and normal & dry wing length of flies subjected to PR50 and PR70 (50% and 70% protein content present in control food respectively) for 20 generations from pre-adult stage. We found that PR fed flies have lower body weight, relative water content (in males), unaltered (PR50%) and higher (PR70%) relative fat content in males, smaller normal and dry body size as compared to control and generations 1 and 2. Interestingly, wing size and pupal size of PR flies are smaller and showed significant effects of diet and generation. Thus, these traits are sex and generation dependent along with an interaction of diet, which is capable of modulating these results variably. Our study suggests that trans-generational effect is more prominent in influencing these traits and wing length might not be a predictor for body size. Taken together, the trans-generational effect of PR on fitness and fitness-related traits might be helpful to understand the underpinning mechanisms of evolution and aging in fruit flies *D. melanogaster*.

**Summary statement:** Twenty generations of protein restricted diet have a diet and generation dependent effect on adult body size, wing length and body weight.

## Introduction

Organisms vary in body size not only across species but also within a particular species. The variations in the body composition can influence phenotypic traits like body size, body weight, etc., while these trait variations can be attributed to various environmental and genetic factors (Robertson, 1960, 1963; de Moed et al., 1997). The environmental factors that can influence organismal body size and weight, including wing length (especially in insects) can be nutrition (Klepsatel et al., 2018), temperature (Nunney and Cheung, 1997; Karan et al., 1998), crowding (Miller and Thomas, 1958; Klepsatel et al., 2018), latitudinal clines (Robinson et al., 2000) and certain cases of laboratory selection pressures for faster development (Yadav and Sharma, 2014), etc. Body size, weight and wing length are certain parameters that ensure the overall fitness of organisms including fruit flies. Thus, variations in these phenotypes can be used to understand the genotypic changes that are bound to occur (Ormerod et al., 2017).

Fruit flies *Drosophila melanogaster* for the past three decades, has been widely used as a model organism for studying aging via nutritional approaches including diet restriction (DR), food dilution, intermittent feeding, etc., (Bass et al., 2007; Grandison et al., 2009; Katewa et al., 2012). The diet of fruit flies commonly comprises of carbohydrates and proteins as the major source, with lipids, vitamins, minerals present in minor quantities. Restricting protein source (yeast) in the fly food is a type of dietary implementation (Protein Restriction; PR henceforth), and is seen to influence a range of fitness and fitness-related traits such as lifespan, fecundity, stress resistance, activity, development time, etc., (Chapman and Partridge, 1996; Katewa et al., 2012; Sisodia and Singh, 2012; Krittika et al., 2019). Interestingly, PR is known to influence traits like body size, weight, and wing length, wherein variations in yeast concentration can significantly alter the body size and weight of the flies, and also have an influence on their wing length in a single generation itself (environment effect; Robertson, 1960, 1963; G*ü*leret al., 2015). This might be due to the sudden change in the protein composition; while it is necessary to assay long term (genetic effect) effect of PR. A high protein diet can yield unaffected pupal size (Reis, 2016), while the long duration of high protein increased body mass (Kristensen et al., 2011), also decreased body weight and fat levels (Morris et al., 2012).

PR from the pre-adult stage is highly debatable as some studies suggest its negative effect on lifespan, body size, fecundity (Robertson, 1960; Hodin and Riddiford, 2000; Udugade et al., 2016); while studies claim its positive effect on lifespan and other traits (Katewa et al., 2012; May et al., 2015; Stefana et al., 2017). The adult body size need not necessarily influence the lifespan of the organism raised under varied nutritional conditions (Tu and Tatar, 2003). Since, nutrition during the pre-adult stage largely determines the size of the adult upon eclosion (Robertson, 1960, 1963; Tu and Tatar, 2003) alongside the influences of juvenile hormones (Mirth et al., 2014), it is essential to study the long-term effect of PR and not just one or two generations (Matzkin et al., 2013). Moreover, assaying for one or two generations might address the immediate effect of parents or grandparents’ diet on the off springs and whether it is maternally inherited or not (Vijendravarma et al., 2010; Valtonen et al., 2012). The current study assays a trans-generational effect of 20 generations (gen 1, 2, and 20) and does not deal with understanding the mode of inheritance (maternal or paternal).

Here we address the effect of 50% and 70% yeast concentrations (as against the control-AL diet) across generations 1, 2 and 20. The PR concentrations of 50% and 70% have been used based on the preliminary PR studies on the lifespan of the flies (unpublished data). This study will address the effect of PR on a single generation, its offspring (generation 2) and also long-term effect (generation 20) of the corresponding protein restriction. The assessed traits include body size, weight, relative water and fat content, pupal size and wing length in the normal and dry conditions of the fly body. After 20 generations of PR implementation, the PR males and females have lower body weight as compared to their control in their respective generations and within the PR generations. Interestingly, the relative water content is higher in females and not males despite the long-term PR diet. Since, the body weight of the DR flies is lower after generations, we found it necessary to assess the relative fat content of the flies. We found that the DR70% males have higher fat storage after 20 generations, while females showed no difference. Moreover, the flies at generation 20 have lower body size as compared to the generations 1 and 2 and their control, showing that the body size and weight might be positively correlated. Since the body size is an adult trait, we measured the intermediate pupal size. The PR50 flies at gen 1 and 2 showed the highest pupal size as compared to PR70 and control, while their size was similar at the end of 20 generations, which might be reflected as a part of smaller adult body size as well. Lastly, since wing length itself can be an indicator of body size (Sokoloff, 1966), measuring the same revealed that PR had a diet and generation dependent effect on producing flies with shorter wings. Thus, after 20 generations, PR diets produce flies with smaller body and wing, lower weight, unaltered pupal size and relative water content (in females). Thus, this study would benefit to understand the influence of diet and/or the genetic effect (generational study) in mediating variations in the assayed traits.

## Materials and methods

### Fly culture and maintenance

The control and DR imposed flies are maintained on 21-day discrete generation cycles with egg collection done exactly on the 21^st^ day of the previous generation cycle. The control flies are fed with AL (*Ad Libitum*) food, while the PR stocks are fed with 50% and 70% yeast as compared to the control (PR50% and PR70%; henceforth). The egg collection for control and PR stocks are done in their respective AL and PR diets (AL diet but with 50% and 70% yeast for PR50 and PR70 stocks respectively). The flies upon eclosion are transferred to plexi-glass cages (25 cm × 20 cm × 15 cm) and are supplemented with their corresponding food. For the following experiments, approximately 30-40 eggs per vial were collected for control and PR and maintained at the temperature of ∼25 °C (±0.5 °C), the humidity of ∼70 %, the light intensity of ∼250 lux in 12:12 hr Light/Dark cycles. The diet manipulations were done only in the yeast concentration present in the control food, wherein we used instant dry yeast from Gloripan.

### Normal body weight and dry weight

We measured the normal and dry body weight of freshly eclosed flies collected in every 2 h intervals. The eggs were collected from the DR stocks over a 2 h window and kept under LD12:12 h. Post eclosion, the virgin male and female flies were separated by anesthetizing with CO_2_. For weighing the normal body weight, flies were weighed post anesthetization using ether (to maintain the flies in the anesthetized state for longer duration), after which the flies were discarded. For the dry body weight assay, the virgin males and females were killed by freezing and were dried for 36 h at 70°C as per the protocol followed elsewhere (Yadav and Sharma, 2014). The normal and dry body weight assay was assessed by weighing a group of 10 males or 10 females per vial, and 5 such vials of randomly chosen flies from the control and the DR stocks were weighed. The body weight of flies was measured using a weighing balance from UniBloc (Shimadzu) AUX220. The relative water content of the flies was calculated by dividing the water content (normal body weight-dry weight) by the normal body weight of the flies as reported elsewhere (Robinson et al., 2000). The relative fat content was assessed by dividing the fat content (dry weight-fat free dry weight) by the dry weight of the flies (Robinson et al., 2000).

### Body size/length and wing length

The protocol for egg collection until the separation of virgin male and female flies for this assay is similar to that followed for body weight assay. The flies’ body size and wing length were measured under a microscope, wherein 30 virgin males and females from the control and DR stocks were assayed. The body size and the wing length of the anesthetized males and females were measured using a microscope from Olympus with a normal ruler (least count 0.5 mm).

## Results

### Normal and dry body weight

To check whether the body size and weight are proportional to each other under the imposed PR, we assayed body weight and size (normal and dry) of the AL and PR stocks at gen 1, 2 and 20 generations of diet imposition. ANOVA followed by post hoc multiple comparisons using Tukey’s test on the normal body weight of the freshly eclosed male and female fruit flies showed a statistically significant effect of diet (D; F_*2,72*_=193.04, p<0.0001), generation (G; F_*2,72*_=99.63, p<0.0001), sex (S; F_*1,72*_=490.70, p<0.0001) and their interaction (D × G× S; F_*4,72*_=9.54, p<0.0001; Table 1a). The results show that PR50 (males) and PR70 (males and females) have significantly lower body weight at gen 20 as compared to their previous generations (Fig. 1A); while PR50 females at gen 20 have lower body weight than gen 2, but not gen 1. The PR50 and PR70 (males and females) have lower body weight at all tested generations against their respective control. Thus, with PR and long-term restrictions, the adult body weight is lower. To assess whether this lower body weight is due to the water content, we weighed the dry weight of the flies.

**Table 1a:**
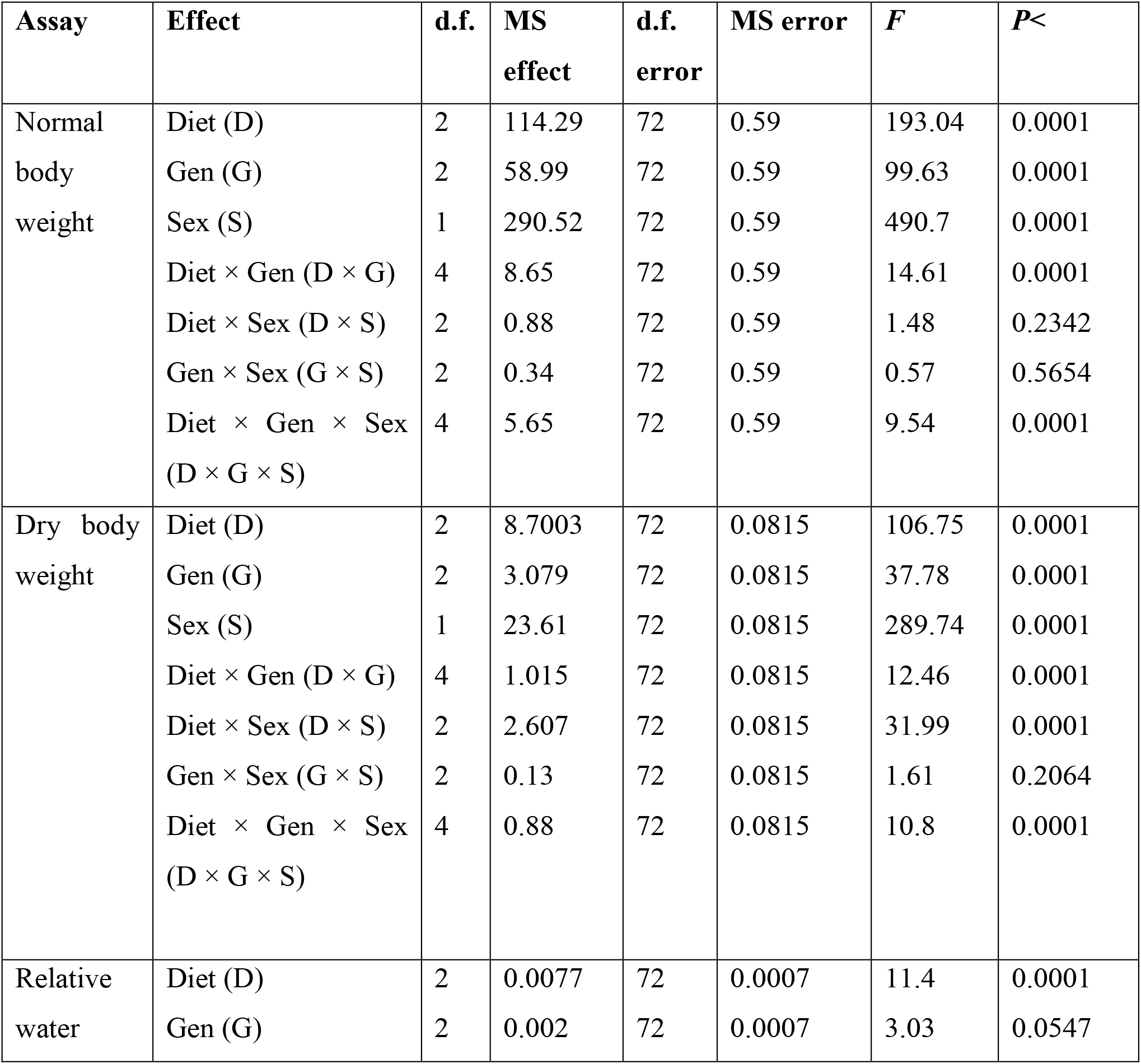
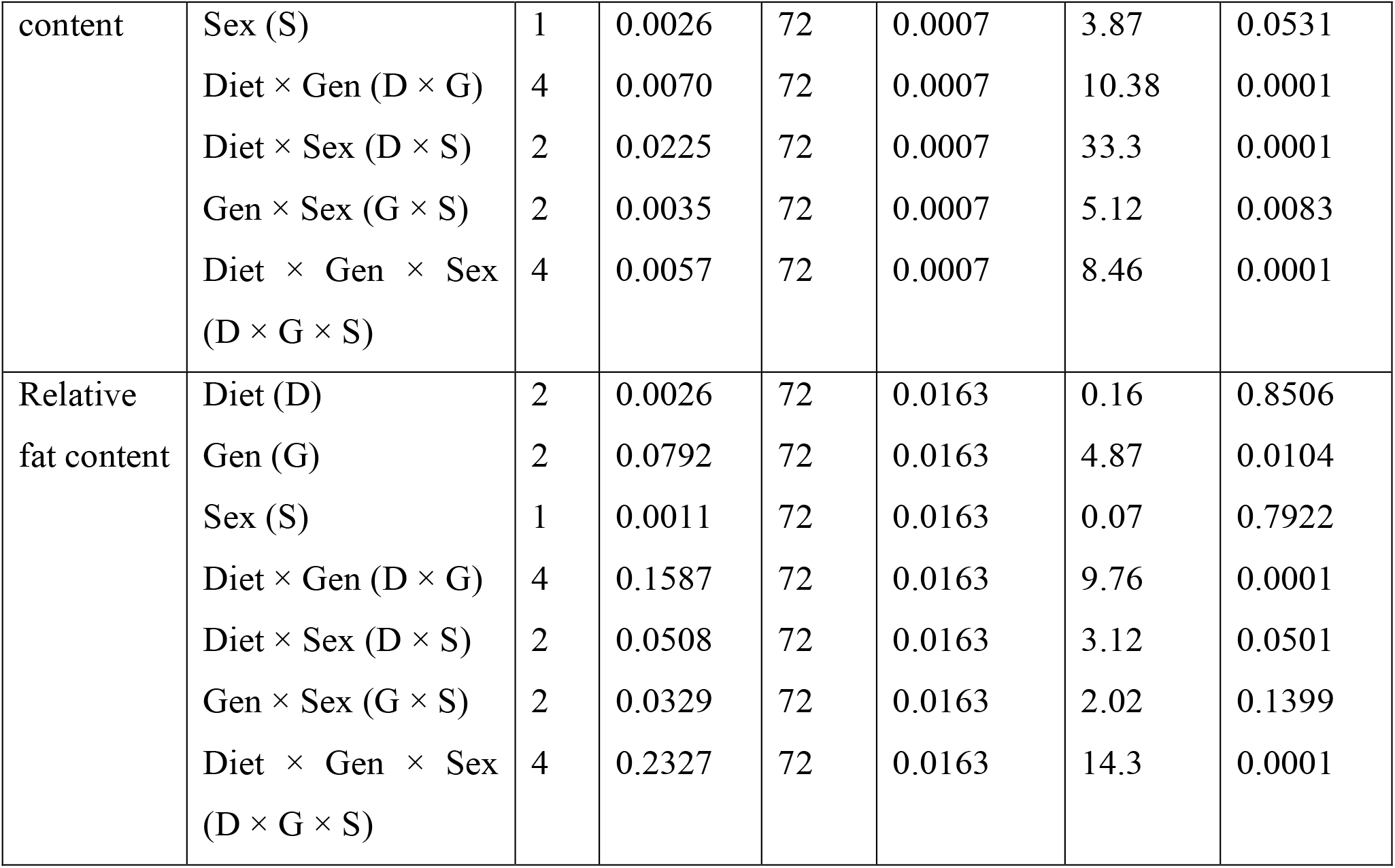
ANOVA details of the normal & dry body weight and relative water & fat content of long-term PR imposed flies.

**Figure 1:**
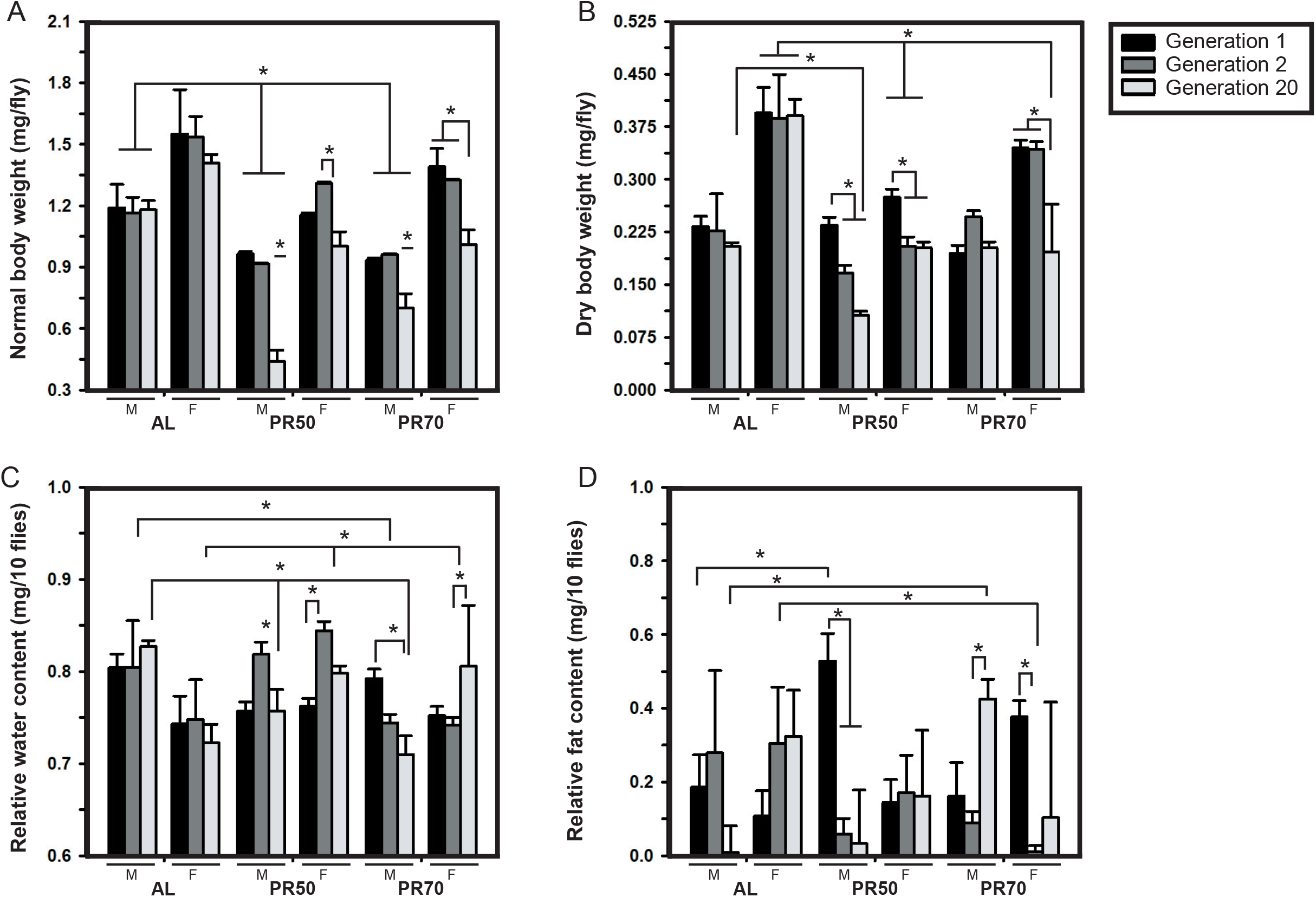
Low weighed males and females under the PR diet for 20 generations. The normal (A) and dry (B) body weight of the flies (varying across generations), shows PR flies weighing lower than that of AL flies at the end of generation 20. The effect of diet on the relative water content (C) is prominent, wherein after 20 generations of PR diets, male and female flies possessed lower and higher water content as compared to AL respectively. The graph represents diet in the *x*-axis and body weight (A, B), relative water content (C), relative fat content (D) in the *y*-axis. The bars and error bars are represented as the mean ± standard deviation (SD). The asterisks on the bars indicate significance levels wherein the *p-*value is <0.05. G1, G2, and G20 represent generation 1, 2, and 20 respectively, while M and F represents males and females respectively.

As expected, the ANOVA on dry body weight of flies revealed statistically significant effect of D (F_*2,72*_=106.75, p<0.0001), G (F_*2,72*_=37.78, p<0.0001), sex (S; F_*1,72*_=289.74, p<0.0001) and their interaction (D × G × S; F_*4,72*_=10.8, p<0.0001; Table 1a). Post hoc multiple comparisons by Tukey’s test revealed significantly decreased dry body weight of PR50 (males and females) at gen 20 have lower dry weight as compared to their gen 1, but not gen 2. Interestingly, PR70 males show no effect of generations as their dry weight is not different, while PR70 females at gen 20 have lower weight as compared to gen 1 and 2 (Fig. 1B). The effect of diet shows that at gen 1, the PR50 females are lower in weight, while the others weigh similar to AL. At gen 2, only PR50 females were lower, but surprisingly at gen 20, PR50 and PR70 (males and females) dry weight difference is prominent and weigh lower than the control. Thus, the results confirm that the body weight of PR flies is lower than the AL due to long term diet implementation.

Since there exist a difference between the normal and dry body weights across generations, we assessed the relative water content in the PR flies. ANOVA on the relative water content revealed statistically significant effect of D (F_*2,72*_=11.4, p<0.0001) and its interaction with generation (D × G; F_*4,72*_=10.38, p<0.0001 and D × G × S; F_*4,72*_=8.46, p<0.0001; Table 1a), but not G (F_*2,72*_=3.03, p<0.0547), sex (S; F_*1,72*_=3.87, p<0.0531). The relative water content of the PR50 flies at gen 2 is comparatively higher than gen 1, while PR70 males and females have higher water content at gen 1 and 20 respectively (Fig. 1C). For the effect of diet across generations, at gen 1, the PR50 and PR70 flies are equal to AL, while at generation 2, the PR50 males have similar water content at that of its respective control, while PR70 males and PR50 females exhibit lower and higher water content respectively. But interestingly at gen 20, PR males have lower relative water content, while in PR females’ it is higher. Thus, long term PR has facilitated higher water content in PR females and not males.

Assessing the direct fat content in flies upon PR can give us information on the fat metabolism in flies. ANOVA on the relative fat content showed statistically significant effect of G (F_*2,72*_=4.87, p<0.0104) and its interaction (D × G; F_*4,72*_=9.76, p<0.0001 and D × G × S; F_*4,72*_=14.3, p<0.0001; Table 1a), but not D (F_*2,72*_=0.16, p<0.8506) and sex (S; F_*1,72*_=0.07, p<0.7922). The post hoc multiple comparisons by Tukey’s test revealed significantly higher fat content in PR70 males at gen 20, while for females it was not significant (Fig. 1D). Interestingly, at gen 2, PR70 females stored lesser fat than the AL females. PR50 flies did not show any difference in their fat content except for PR50 males which showed higher fat than AL in gen 1 (Fig. 1D). Therefore, with respect to the fat content, long term PR has facilitated higher fat content in PR70 males alone after 20 generations, while no significant change was observed in females. This reiterates that the fat accumulation might be sex-dependent and diet dependent as well, given that trans-generational effect is put into consideration.

### Normal and dry body size

The flies maintained on PR50% and 70% for 20 generations from the pre-adult stage were measured for their normal and dry body size. ANOVA on the normal body size of freshly eclosed adult males and females showed a statistically significant effect of diet (D; F_*2,522*_=38.51, p<0.0001), generation (G; F_*2,522*_=98.19, p<0.0001), sex (S; F_*1,522*_=611.60, p<0.0001) and their interaction (D × G; F_*4,522*_=21.79, p<0.0001 and D × S; F_*2,522*_=17.52, p<0.0001) (Table 1b; Fig. 2A), but not D × G × S (F_*4,522*_=2.07, p<0.0830). Further, post hoc multiple comparisons using Tukey’s HSD test revealed the generation effect is prominent in the PR flies as their body size at gen 20 is comparatively smaller than the previously tested generations (1 and 2). The interaction between all three tested factors is not significant probably due to the fact that body size is capable of higher perturbation to environmental factors. The effect of diet shows that at gen 1, PR males do not show any difference in body size, while PR females are smaller. But surprisingly at gen 2, the PR fed males are larger than AL, while females are similar in size as that of AL, while at gen 20, PR males and females are smaller than the AL flies. Thus, after 20 generations the PR produces smaller flies as compared to AL even though minor fluctuations in their body size were observed at gen 1 and 2.

**Table 1b:** ANOVA details of the normal and dry body size of long-term PR imposed flies.

| Assay | Effect | d.f. | MS<br>effect | d.f.<br>error | MS error | *F* | *P*< |
| --- | --- | --- | --- | --- | --- | --- | --- |
| Normal<br>body size | Diet (D) | 2 | 1.036 | 522 | 0.027 | 38.51 | 0.0001 |
|  | Gen (G) | 2 | 2.641 | 522 | 0.027 | 98.19 | 0.0001 |
|  | Sex (S) | 1 | 16.45 | 522 | 0.027 | 611.60 | 0.0001 |
| | Diet × Gen (D × G) | 4 | 0.586 | 522 | 0.027 | 21.79 | 0.0001 |
| | Diet × Sex (D × S) | 2 | 0.471 | 522 | 0.027 | 17.52 | 0.0001 |
| | Gen × Sex (G × S) | 2 | 0.143 | 522 | 0.027 | 5.32 | 0.0052 |
| | Diet × Gen × Sex<br>(D × G × S) | 4 | 0.056 | 522 | 0.027 | 2.07 | 0.083 |
| Dry body<br>size | Diet (D) | 2 | 0.374 | 522 | 0.018 | 21.23 | 0.0001 |
|  | Gen (G) | 2 | 2.870 | 522 | 0.018 | 163.13 | 0.0001 |
|  | Sex (S) | 1 | 6.394 | 522 | 0.018 | 363.37 | 0.0001 |
| | Diet × Gen (D × G) | 4 | 0.724 | 522 | 0.018 | 41.12 | 0.0001 |
| | Diet × Sex (D × S) | 2 | 0.025 | 522 | 0.018 | 1.42 | 0.2438 |
| | Gen × Sex (G × S) | 2 | 0.0004 | 522 | 0.018 | 0.03 | 0.9729 |
| | Diet × Gen × Sex<br>(D × G × S) | 4 | 0.015 | 522 | 0.018 | 0.86 | 0.4907 |

**Figure 2:**
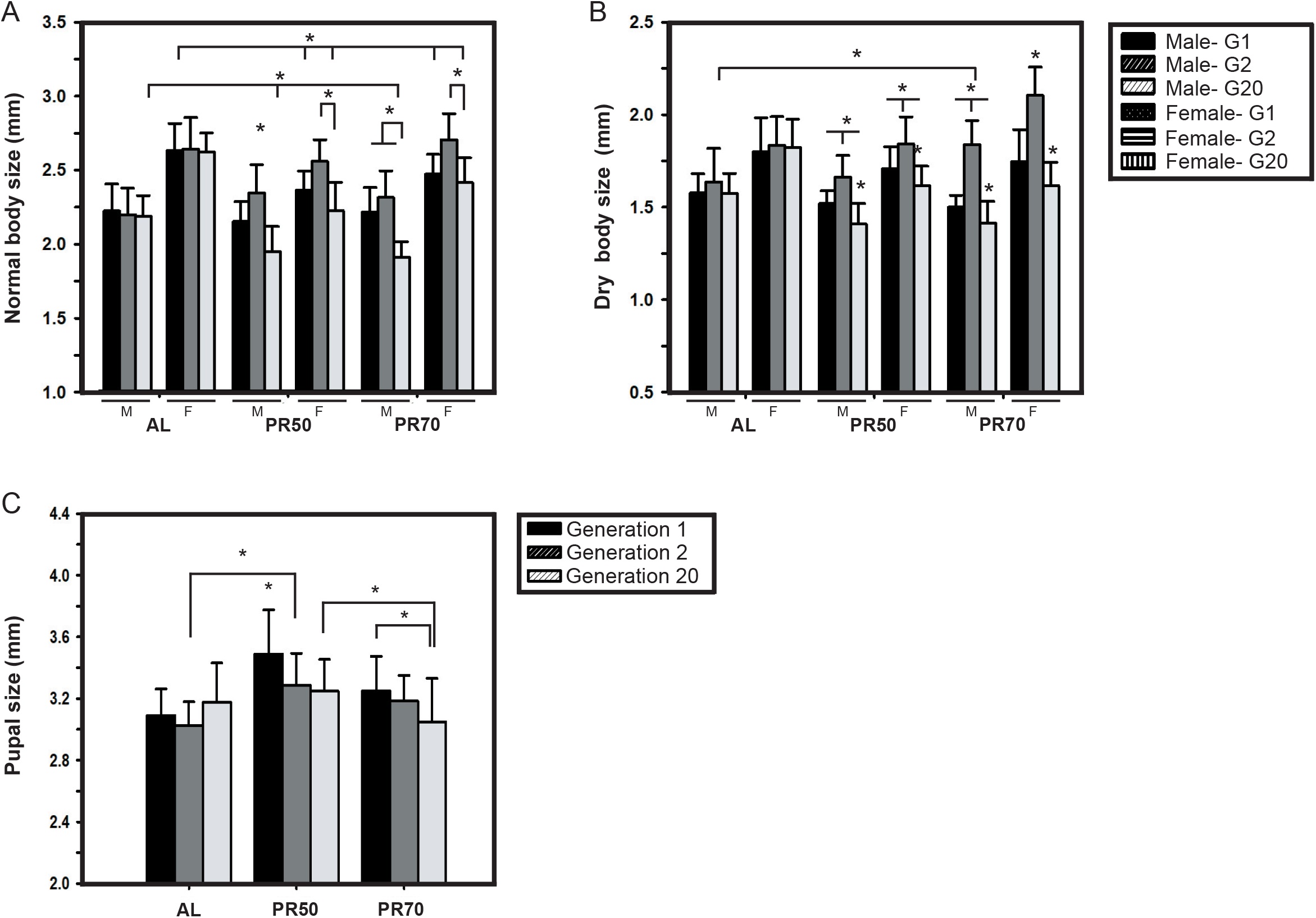
Smaller flies and unaltered pupal size due to the PR diet for 20 generations. The effect of diet and generation on the normal (A) and dry (B) body size of the flies are variable, wherein the normal body size of PR flies is lower than their control at the end of the 20 generations. The pupal size (C) of PR50 flies were the highest at gen 1 as compared to PR70 and control, but after 20 generations were similar to AL. The graph represents diet in the *x*-axis and body size (A, B), pupal size (C) in the *y*-axis. All other details are the same as in Figure 1.

Further, ANOVA followed by multiple comparisons by Tukey’s HSD test on the dry body size showed statistically significant effect of D (F_*2,522*_=21.23, p<0.0001), G (F_*2,522*_=163.13, p<0.0001), S (F_*1,522*_=363.37, p<0.0001) and their interaction (D × G; F_*4,522*_=41.12, p<0.0001; Table 1b; Fig. 2B), but not D × S (F_*2,522*_=1.42, p<0.2438) and D × G × S (F_*4,522*_=0.86, p<0.4907). All the PR flies at gen 2 are bigger as compared to gen 1 and 20, except for PR70 males wherein their dry body size is similar to that observed at gen 1. The effect of diet on the dry body size revealed that PR flies are similar in size to AL at gen 1, while at gen 2 the PR70 (males and females) are bigger than AL. Similar to the results of normal body size, the PR flies are smaller than their control flies at gen 20 and might to due to the factors discussed earlier with normal body size. Surprisingly, there exist changes in the response of PR diet on the normal and dry body size, showing that the normal body size and dry body size might not be equivalent and the difference between them is not constant, and the reason might be attributed to the various forms of storage reserves.

### Pupal size

ANOVA followed by Tukey’s HSD test on the normal wing length showed a statistically significant effect of D (F_*2,261*_=29.34, p<0.0001), G (F_*2,261*_=7.96, p<0.0004) and their interaction (D × G; F_*4,261*_=5.91, p<0.0001; Fig. 2C; Table 1c). Post hoc multiple comparisons by Tukey’s test showed that among PR50 flies across generations, gen 1 was the highest, while at gen 2 and 20 they were similar. Across diets, PR50 flies had a higher pupal size in gen 1, 2 and gen 20 as compared to AL and PR70 flies. Thus, showing that the PR50 flies have pupal size higher as compared to the control in all the generations, but within its generations, the observed highest pupal size at generation 1 might have been a startle response for PR.

**Table 1c:** ANOVA details of the pupal size of long-term PR imposed flies.

| Assay | Effect | d.f. | MS<br>effect | d.f.<br>error | MS error | *F* | *P*< |
| --- | --- | --- | --- | --- | --- | --- | --- |
| Pupal size | Diet (D) | 2 | 1.452 | 261 | 0.049 | 29.34 | 0.0001 |
|  | Gen (G) | 2 | 0.394 | 261 | 0.049 | 7.96 | 0.0004 |
| | Diet × Gen (D × G) | 4 | 0.292 | 261 | 0.049 | 5.91 | 0.0001 |

### Normal and dry wing length

Post body size assessments, we intended to assay the wing length as it is commonly thought to be a measure of body size as mentioned earlier. ANOVA on the normal wing length showed statistically significant effect of D (F_*2,522*_=72.28, p<0.0001), G (F_*2,522*_=28.07, p<0.0001), S (F_*1,522*_=301.92, p<0.0001) and their interaction (D × G; F_*4,522*_=10.6, p<0.0001; and D × G × S; F_*4,522*_=4.6, p<0.0012; Table 1d). Interestingly, multiple comparisons by Tukey’s test within diets across generations shows that gen 20 females have wing length similar to gen 2, while the males have lower wing length compared to gen 2. Moreover, PR50 females have smaller wing length as compared to AL in all tested generations; while PR70 females have smaller wings as compared to AL in the first generation alone (Fig. 3A). The effect of diet shows that PR flies (males and females) have shorter wings than AL flies at gen 20. Thus, the concept of wing length as a measure for body size might not hold in the presence of dietary parameters influencing them across generations.

**Table 1d:**
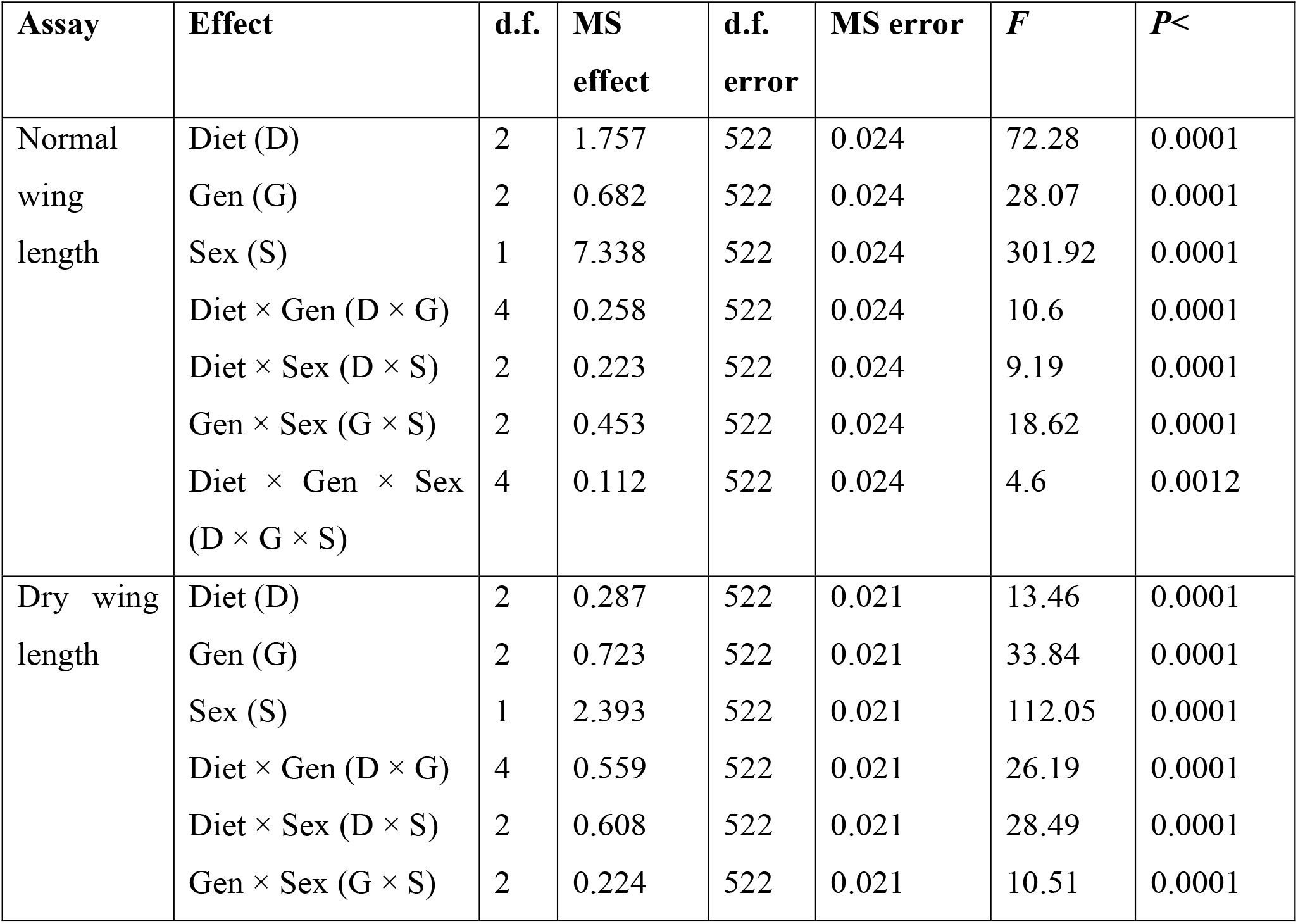
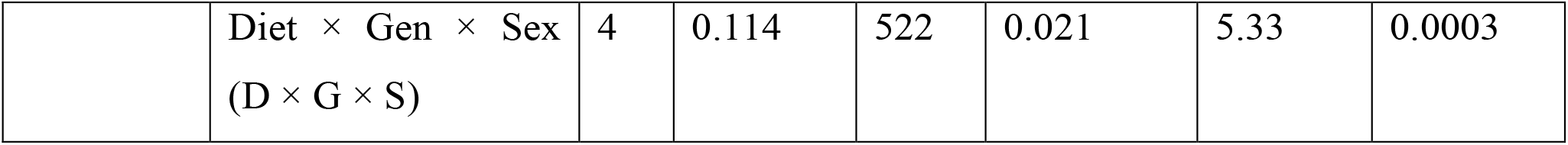
ANOVA details of the normal and dry wing length of long-term PR imposed flies.

**Figure 3:**
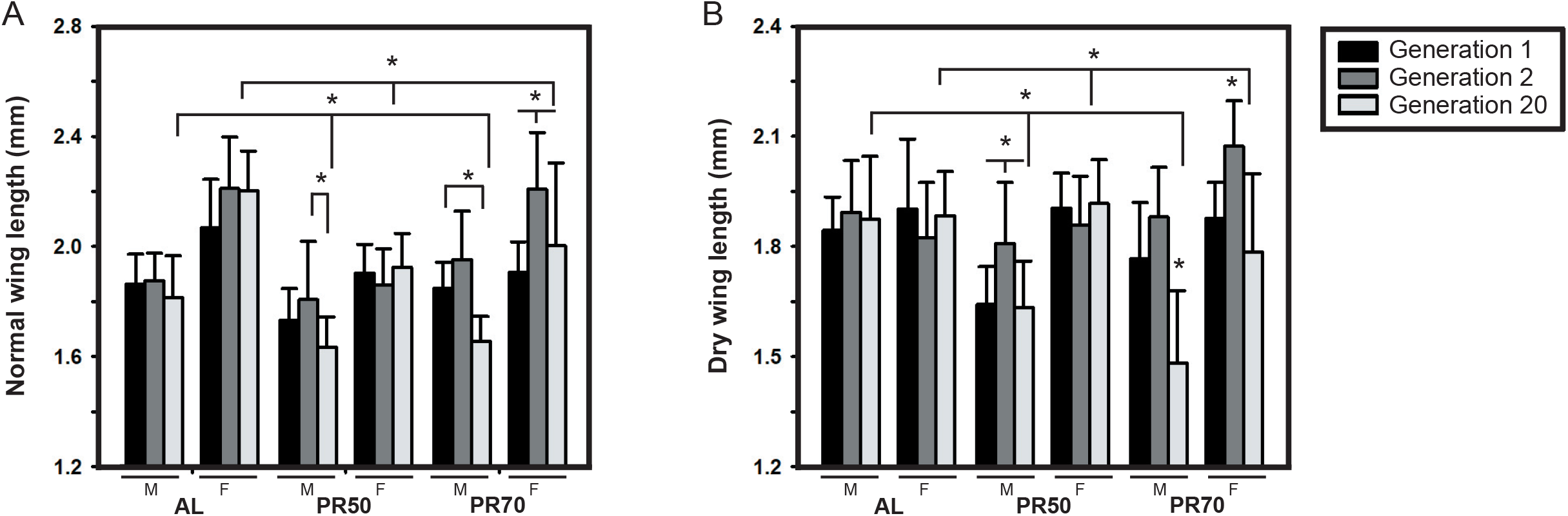
Flies wing length reduced due to long-term PR diet imposition. The normal (A) and dry wing length (B) of the PR flies are smaller than the AL flies post 20 generations, but the pattern of variations across generation is different from that witnessed for body size. The graph represents diet in the *x*-axis and wing length on the *y*-axis. All other details are the same as in Figure 1.

ANOVA on dry wing length of fruit flies revealed significant effect of D (F_*2,522*_=13.46, p<0.0001), G (F_*2,522*_=33.84, p<0.0001), S (F_*1,522*_=112.05, p<0.0001) and their interaction (D × G; F_*4,522*_=26.19, p<0.0001 and D × G × S; F_*4,522*_=5.33, p<0.0003; Table 1d). Multiple comparisons of dry wing length by Tukey’s test revealed results similar to the normal wing length wherein PR flies at gen 20 had significantly smaller wings than the control (Fig. 3B). Thus, even though PR yield flies with shorter wings at the end of 20 generations (similar to dry body size), it is not true across generations, while it explicitly shows significant interaction between diet, sex and generations.

## Discussion

### Normal and dry body weight

Our results are similar to the results of another study (Vijendravarma et al., 2010) that show the effect of diet (parental) on different traits (including body weight) of fruit flies, and it also suggests that these observed differences might be due to the maternal effects and the long-term DR imposition. Further, we did not expect variations in the AL body weight across generations, and convincingly their body weight and relative water content were unaffected, thereby providing convincing results for the control flies. In the PR50 and 70% flies, there might be due to the effect of parental diet on the normal body weight of the flies as suggested elsewhere (Vijendravarma et al., 2010; Valtonen et al., 2012), because the PR flies showed lower body weight at the end of 20 generations. Moreover, the lower body weight of PR flies after 20 generations can be thought to be in line with the study of Kristensen et al., 2011, which reported the protein-rich diet for 17 generations yielded bigger and thereby fatter flies as compared to the control. But surprisingly, PR70 males have higher fat content than AL at the end of 20 generations, which makes it contrary to Kristensen et al., 2011, even though the study reported the implementation of protein-rich diet. The difference in relative fat content upon sex-based effect is similar to that reported elsewhere, which showed fat content varied with significant effect of sex (Kristensen et al., 2011) and density of larvae (Zwaan et al., 1991; Kristensen et al., 2011), while relative fat content varied with sex alone (Kristensen et al., 2011). But the present study is contrary to the same clinal study in the fact that our male flies have lower relative water content and higher fat content at the end of 20 generations. Interestingly, the dry weight of PR males is more stable than the females and is in line with the study reported elsewhere (Karan et al., 1998). Thus, long term PR implementation suggests the existence of a plastic response to diet as compared to the genetic effect in case of dry weight and sex (Kristensen et al., 2011).

### Body size and Pupal size

The body size of PR flies is smaller at generation 20 as compared to their previous generations and control. These results are contrary to the study of Chippindale et al., 1996, which reported that bigger adult body size is associated with increased fitness of the flies. Since the fitness of the organism is assessed based on its reproductive capacity and the ability to withstand stress, our results might have a positive effect despite a smaller body size. Moreover, populations of *D. melanogaster* selected for faster development (FD) exhibit reduced body size, lower lifespan and reduced fitness (Yadav and Sharma, 2013, 2014), but the PR flies show higher lifespan in spite of their reduced body size (unpublished data). Surprisingly, flies with large body size exhibit lower larval viability even though they appear to contribute to the adult fitness (Partridge and Fowler, 1993). Since the females that mated with smaller males appeared more fecund and also copulated longer (Pitnick, 1991), the duration of copulation and offspring number are dependent on the female body size and inversely related to the male body size (Pitnick, 1991; Lefranc and Bundgaard, 2000). In tandem with these studies, even though the large females differ in size as compared to the smaller females, does not guarantee significant difference in their ovariole number (Lefranc and Bundgaard, 2000), even though yeast restriction reduces ovariole number (Tu and Tatar, 2003) and high protein diet is known to increase the same with a possible trade-off in the egg to adult viability in *D. ananassae* (Sisodia and Singh, 2012). Moreover, since the males of *D. melanogaster* prefer smaller females for first mating and then undergo adaptive discrimination (Byrne and Rice, 2006) or plasticity for mate selection by males (Edward and Chapman, 2013), we can conclude that body size may be one of the many traits that are assessed to choose a potential mate but not a primary one. Hence, the smaller body size of the PR flies might not be a threat for its mate choice, reproductive success or larval viability in our study, even though the fecundity of our flies remains to be tested.

The pupal size at PR50 flies recorded the highest size as compared to the control and PR70 flies, while gen 1 flies of PR50 yielded highest pupal size compared to gen 20 (Fig. 2C). Since high protein diet did not confer any change in the pupal size of the flies, but high carbohydrate diet resulted in smaller pupa (Reis, 2016), it is surprising to see pupal size difference upon protein restriction. This is in line with the results of Deas et al., 2019; that suggests more susceptibility of pupal mass change in poor diet than that of rich nutrient diet, in addition to exhibiting effects of parental and grandparental diet (Deas et al., 2019). Overall, our results also show that diet and generation have a differential role of different traits as suggested elsewhere (Deas et al., 2019).

### Normal and dry wing length

The concentrations of nutrients (yeast and sugar) in the fly diet play an important role in the wing length of females than in males (Güler et al., 2015). In our study, since the concentration of sugar was kept constant, the observed variations in wing length show that the yeast alone can modulate this trait. The PR flies showed smaller wings after 20 generations depicting generational effect while there exists variable results due to sex difference. This is contrary to Güler et al., 2015, where female’s wing length varies with yeast manipulations while males vary with sugar level variations. There are various other factors capable of modulating wing length like temperature and latitudinal clines (David et al., 1983; de Moed et al., 1997), wherein care was taken to avoid such temperature perturbations. It is also seen that wing length (similar to that of body size) can be influenced by altitude in *Drosophila* species (Stalker and Carson, 1948; Tantawy, 1965). There exists a difference in the PR and generation effect on the trend of body size and wing length variations and probably, is in contrast to the study of Sokoloff, 1966, which stated that wing length can serve as a parameter for estimating fly’s body size.

## Conclusion

The results of our study reports that post 20 generations, the PR flies tend to exhibit lowered body size, weight, wing length and pupal size which is highly dependent on diet, sex and generation. It is evident that wing length cannot be an accurate measure of body size and so does the concept of bigger flies are larger, due to perturbations in body reserves of water and fat. Therefore, this study along with previous studies of PR can be taken to suggest that diet, sex and generational effects are capable of interfering with the phenotypic traits of the flies, and thus the genetic and environment effect is highly prominent.

## Acknowledgments

We thank Dr. Usha Nagarajan, Fly Lab, SASTRA Deemed to be University for the microscope facility for measuring body size and wing length. We thank N. Ramesh (SASTRA University) for helping us with population maintenance.

## Author contributions

S. K. & P. Y. conceived and designed the experiments; S. K. performed the experiments, and analyzed the data. S. K. & P. Y. wrote initial versions of the manuscript, corrected, read and approved the final version of the manuscript.

